# Inhibition of the Ventral Hippocampus Increases Alcohol Drinking

**DOI:** 10.1101/786178

**Authors:** William C. Griffin, Jennifer A. Rinker, John J. Woodward, Patrick J. Mulholland, Howard C. Becker

**Affiliations:** Charleston Alcohol Research Center, Department of Psychiatry and Behavioral Science; Department of Neuroscience, Medical University of South Carolina; Ralph H. Johnson VA Medical Center, Charleston, SC 29425-0742

## Abstract

The function of the ventral hippocampus (vHC) supports many behaviors, including those related to reward seeking behaviors and drug addiction. We used a mouse model of alcohol dependence and relapse to investigate the role of the vHC in alcohol (ethanol) drinking. One experiment used a chemogenetic approach to inhibit vHC function while ethanol dependent and non-dependent mice had access to ethanol. Interestingly, the non-dependent mice expressing an inhibitory DREADD in the VHC showed a significant increase in ethanol drinking (∼30%) after hM4Di DREADD activation with clozapine-n-oxide (CNO; 3 mg/kg, ip.) compared to vehicle. On the other hand, ethanol dependent mice, which were already drinking significantly more ethanol than non-dependent mice, only had a slight, non-significant increase in drinking after CNO challenge. In a separate group of dependent and non-dependent mice, GCaMP6f calcium-dependent activity was recorded in the vHC while mice were actively drinking ethanol. These data showed that once mice were rendered ethanol dependent and were drinking more ethanol than the non-dependent mice, calcium signaling in the ventral hippocampus decreased (∼45%) in the ethanol dependent mice compared to the non-dependent mice. Together, these findings suggest that ethanol dependence reduces activity of the ventral hippocampus and that reduced function of this brain region contributes to increased ethanol drinking.

## Introduction

The consequences of heavy alcohol drinking extract significant personal, medical and economic costs (Rehm *et al.* 2014; Rehm *et al.* 2017). Unfortunately, neurobiological adaptations that perpetuate harmful drinking are not fully understood and animal models remain important for the basic discovery of mechanisms and treatments of alcohol use disorder. Mice rendered alcohol (ethanol) dependent display significant escalations of ethanol drinking as compared to non-dependent animals (Becker 2013; Becker & Lopez 2004; Griffin 2014; Griffin *et al.* 2014; Griffin *et al.* 2015; Griffin *et al.* 2009b; Lopez & Becker 2005). In these studies, C57BL/6J (B6) mice are exposed to repeated cycles of chronic intermittent ethanol (**CIE**) vapor exposure which are interleaved with periods of voluntary drinking. Under these conditions, CIE exposed mice increase ethanol consumption compared to air-exposed control mice (**CTL**). Similar findings have been reported by other laboratories in mice (DePoy *et al.* 2013; Dhaher *et al.* 2008; Finn *et al.* 2007; Jeanes *et al.* 2011) and rats (Gilpin *et al.* 2008; Hansson *et al.* 2008; O’Dell *et al.* 2004; Roberts *et al.* 1996). The escalated drinking observed in ethanol dependence is likely due to adaptations in specific neurobiological systems regulating ethanol intake.

It is well known that the hippocampus plays a prominent role in mediating spatial orientation, navigation and learning and memory. Interestingly, preclinical lesion and inactivation studies indicate that the more anterior and dorsal areas of the hippocampus have the largest role in spatial orienting and learning behaviors, while the posterior and ventral areas of the hippocampus, though still contributory to spatial navigation, appear to also participate in other realms of behavior (Bannerman *et al.* 2004; Fanselow & Dong 2010; Kanoski & Grill 2015; Pennartz *et al.* 2011; Strange *et al.* 2014). Nevertheless, it has been shown that increased activity of the ventral hippocampus promotes conditioned place preference for social reward (LeGates *et al.* 2018) while inactivation diminishes context-mediated drug (Bossert *et al.* 2016; Lasseter *et al.* 2010; Rogers & See 2007; Sun & Rebec 2003) and alcohol reinstatement (Marchant *et al.* 2016). Thus, the ventral hippocampus processes salient contextual information.

Interestingly, some reports indicate that the vHC plays an important role in motivation and reward that extends beyond simply processing external contextual cues. For example, studies indicate that lesion or reversible inactivation of the vHC increases food reward (Ferbinteanu & McDonald 2001), increases feeding behavior in novel situations (Bannerman *et al.* 2002a; Bannerman *et al.* 2002b), decreases fear and anxiety (Kjelstrup *et al.* 2002; Maren & Holt 2004). Recent data also suggests a role for the ventral hippocampus in regulating habit and goal motivated behaviors (Barker *et al.* 2019) as well as increases in exploration of aversive contexts in approach-avoidance studies (Ito & Lee 2016; Schumacher *et al.* 2016). Collectively, these reports indicate that reduced ventral hippocampus function can facilitate behavior, suggesting the ventral portion of the hippocampus participates in the active inhibition of at least some behaviors (Schwarting & Busse 2017). Thus, the role of the ventral hippocampus in motivation and reward is complex and in need of further study. Given the broad array of behaviors influenced by the ventral hippocampus, it is reasonable to predict that the ventral hippocampus plays an important role in regulating ethanol drinking. Studies in this report address the hypothesis that alcohol dependence reduces ventral hippocampal function and, as a consequence, reduced ventral hippocampal activity promotes increased ethanol consumption.

## Methods

### Subjects

Male C57BL/6J mice (10 weeks of age) were obtained from Jackson Laboratories (Bar Harbor, ME), individually housed and maintained in an AAALAC accredited animal facility under a 12 hr light cycle. Mice had free access to food and water at all times. All experimental protocols were approved by the Institutional Animal Care and Use Committee at the Medical University of South Carolina and were consistent with the guidelines of the NIH Guide for the Care and Use of Laboratory Animals.

Generally, Mice were given access to ethanol using limited access procedures (described below). After establishing stable ethanol intake, mice were separated into chronic intermittent ethanol exposure (**CIE**; ethanol dependent) and control (**CTL**; non-dependent) groups. The EtOH group received chronic intermittent exposure to ethanol vapor in inhalation chambers (16 hr/day for four days) while CTL mice were similarly handled but received air exposure (see below). Limited access drinking sessions were suspended during inhalation exposure. After CIE exposure, mice entered a 72 hr abstinence period and then were tested for voluntary ethanol intake for five consecutive days using limited access conditions as before. Cycles of CIE exposure interspersed with Test weeks of voluntary drinking were repeated several times for both experiments.

### Stereotaxic Surgery

Mice were anesthetized using isoflurane mixed with medical-grade air (4% induction, 2% maintenance), given a subcutaneous injection of 5 mg/kg carprofen (Pfizer, Inc), and placed in a Kopf stereotaxic instrument with digital display (Model 942). For Experiment 1, mice received bilateral infusion of either a virus containing an inhibitory DREADD (AAV(8)-CaMKIIa-hM4Di-mCherry; Addgene, Inc.) or a control virus expressing only mCherry (AAV(8)-CaMKIIa-mCherry; Addgene, Inc.) into the vHC [relative to bregma, AP: −3.4 mm, ML: ±3.0 mm, DV: −4.75mm], relative to Bregma (Franklin & Paxinos 2008). For Experiment 2, mice received a unilateral infusion of a virus to drive expression of a geneitically encoded calcium indicator GCaMP6f (AAV-CamKIIa-GCaMP6f-WPRE; Addgene, Inc) followed by positioning of a fiber implant (MFC_400/430-0.48_5.3mm_MF2.5_FLT; Doric Lenses, Inc) over the viral infusion site (fiber lowered to DV −4.65) which was secured in place by light-cured dental resin (Griffin *et al.* 2014).

### Limited Access Drinking Procedures

Mice were trained to drink ethanol in the home cage as previously described (Becker & Lopez 2004; Griffin 2014; Griffin *et al.* 2009a; Lopez & Becker 2005), with the exception that both experiments started with mice drinking 15% (v/v) ethanol without the sucrose fading procedure used before. The 2-hr drinking sessions started 3 hours into the dark phase Monday through Friday. The amount of ethanol consumed was determined by weighing bottles before and after the access period, spillage was accounted for by measuring bottle weights maintained on empty cages. Mouse body weights were recorded weekly. There were some differences in the limited access procedures between Experiment 1 and 2, noted below in the sections describing them.

### Chronic Intermittent Ethanol (CIE) Exposure Procedures

Chronic intermittent ethanol vapor (or air) exposure was delivered in Plexiglas inhalation chambers as previously described (Becker & Lopez 2004; Griffin 2014; Griffin *et al.* 2009a; Lopez & Becker 2005). Chamber ethanol concentrations were monitored daily and air flow was adjusted to maintain ethanol concentrations within a range that yielded stable blood ethanol levels throughout exposure (175-225 mg/dl). Mice were placed in inhalation chambers at 1600 hr and removed 16-hr later at 0800 hr. Before each 16-hr exposure during CIE treatment, EtOH mice were administered ethanol (1.6 g/kg; 8% w/v) and the alcohol dehydrogenase inhibitor pyrazole (1 mmol/kg) to maintain a stable level of intoxication during each cycle of ethanol vapor exposure (Griffin *et al*, 2009a). CTL mice were handled similarly but received injections of saline and pyrazole. All injections for these procedures were given intraperitoneally in a volume of 20 ml/kg body weight. These procedures were the same for Experiments 1 and 2.

### Experiment 1: Gi-DREADD Activation

During Experiment 1, mice had access to both ethanol and tap water, as an alternative fluid, and the position of ethanol and water bottles were alternated randomly to avoid side preferences. In this study, mice were also habituated to daily intraperitoneal (ip.) injections of normal saline (10 ml/kg) given 30 min prior to ethanol access. Finally, during the 4^th^ Test week, mice were challenged with Clozapine-N-Oxide (CNO; Tocris, Inc) dissolved in saline (0.9%) in a cross-over design such that all mice were tested with vehicle and one dose of CNO (3mg/kg; ip.) to examine changes in ethanol consumption when Gi-DREADDs were activated by CNO in the ventral hippocampus.

### Experiment 2: Fiber Photometry Procedures

For the fiber photometry in Experiment 2, mice had access to only a single bottle of ethanol during the 2-hr access period, as described previously (Braunscheidel *et al.* 2019; Rinker *et al.* 2018). Additionally, the fiber photometry procedures required that mice were tethered during acquisition of GCaMP6f transients and that lickometers (Med Associates, Inc) are used to capture bottle contact data in real time in order to time-lock licking/consummatory behavior with GCaMP6f activity. Immediately after surgery, mice were acclimated to living around-the-clock in lickometer cages which included *ad libitum* access to water from bottles connected to the lickometer system and *ad libitum* food available on the floor of the cage. After several days, mice were connected to a dummy tether each day at the beginning of the dark phase and remained tethered for approximately 6 hours. After 1-2 weeks, ethanol was provided 3 hours into the dark phase and baseline drinking established followed the CIE exposure procedures described above.

Photometry data were acquired using acquisition equipment based on that described by the Deisseroth lab with modifications (Braunscheidel *et al.* 2019). Illumination was provided by 405 nm and 470 nm fiber collimated LEDs (Thorlabs; 30 µW per channel) connected to a four-port fluorescence mini-cube (Doric Lenses, Inc). The combined LED output passed through a 400 µm optical fiber (0.48 NA) pigtailed to a rotary joint (FRJ_1×1_PT; Doric Lenses, Inc) that was housed in a custom-built balance arm. The patchcord connected to the implanted fiber using a pinch connector (ADAF2; Thorlabs, Inc). Emission light was focused onto a photodetector (Newport model 2151; DC low setting) low-passed filtered at 3 Hz and sampled at 6.1 kHz by a RZ5P photometry multiprocessor (TDT, Inc) controlled by Synapse software (TDT, Inc). Excitation light was sinusoidally modulated at 531 Hz (405 nm) and 211 Hz (470 nm) via software control of an LED light driver (Thorlabs). Real-time demodulated emission signals from the two channels were acquired at a frequency of 1017.253 Hz and stored offline for analysis. While the mice were drinking, the lickometer system generated TTLs that were sent to the multiprocessor, signaling contacts at the ethanol bottle.

### Photometry Data Analysis

Raw data collected from the photoreceiver were low-pass filtered and then the extracted 470 nm and 405 nm signals were normalized and analyzed using custom-written scripts in MATLAB (Mathworks, Inc.), following previously published methods (Braunscheidel *et al.* 2019; Rinker *et al.* 2018). To normalize for differing rates of photobleaching (slope measured over single recordings sessions, mean ± sem; 405 nm −0.056 ± 0.001, 470 nm −0.0042 ± 0.0004, N=45), the 405 nm and 470 nm signals for each channel were first fitted to a polynomial versus time. The Ca^++^-independent isosbestic 405 nm signal was then subtracted from the Ca^++^-dependent 470 nm signal, to remove movement artifacts and calculate the final Δ*F/F* time series. Additionally, in these recordings, the mean of the negative values was first subtracted from the normalized Δ*F*/*F* to establish a new baseline of the recordings for comparing changes during ethanol consumption. A licking bout for ethanol was defined as >6 Hz licking rate lasting ≥500 ms. The calcium transients were time-locked to bouts, and the GCaMP6f data before, during and after each bout were extracted.

### Statistical Analysis

Analysis was conducted in SPSS Version 25. Dependent variables were analyzed by factorial analysis of variance (ANOVA) with repeated measures as necessary with *p<0.05* as the criteria for significance. For post-hoc analyses, multiple comparisons Sidak’s test as indicated.

## Results

### Experiment 1

#### Ethanol Drinking Prior to CNO Challenge

In this study, mice underwent 3 cycles of CIE exposure prior to challenge with CNO and these data are summarized in **Figure 1A**. As expected from prior work (Griffin *et al.* 2014; Griffin *et al.* 2015; Griffin *et al.* 2009b), the ethanol dependent (CIE) mice increased ethanol intake significantly over time compared to the relatively stable drinking the CTL group. Further, there was no effect of virus type (mCherry vs hM4Di) on ethanol drinking in either group. This assessment was supported by 3-way ANOVA (2 Groups) X (2 Viruses) X (4 Time points) with Time used as a repeated measure. The 3-way interaction was not-significant [F (3,102) >0.05] and neither was the Time X Virus interaction [F (3,102) >0.05]. However, the Group X Time interaction was significant [F (3,102) <0.05]. Post-hoc analysis of these data show that the CIE mice consumed more ethanol in the Test drinking weeks compared to the CTL group (*p<0.05), as well as their own baseline (^p<0.05).

**Figure 1.**
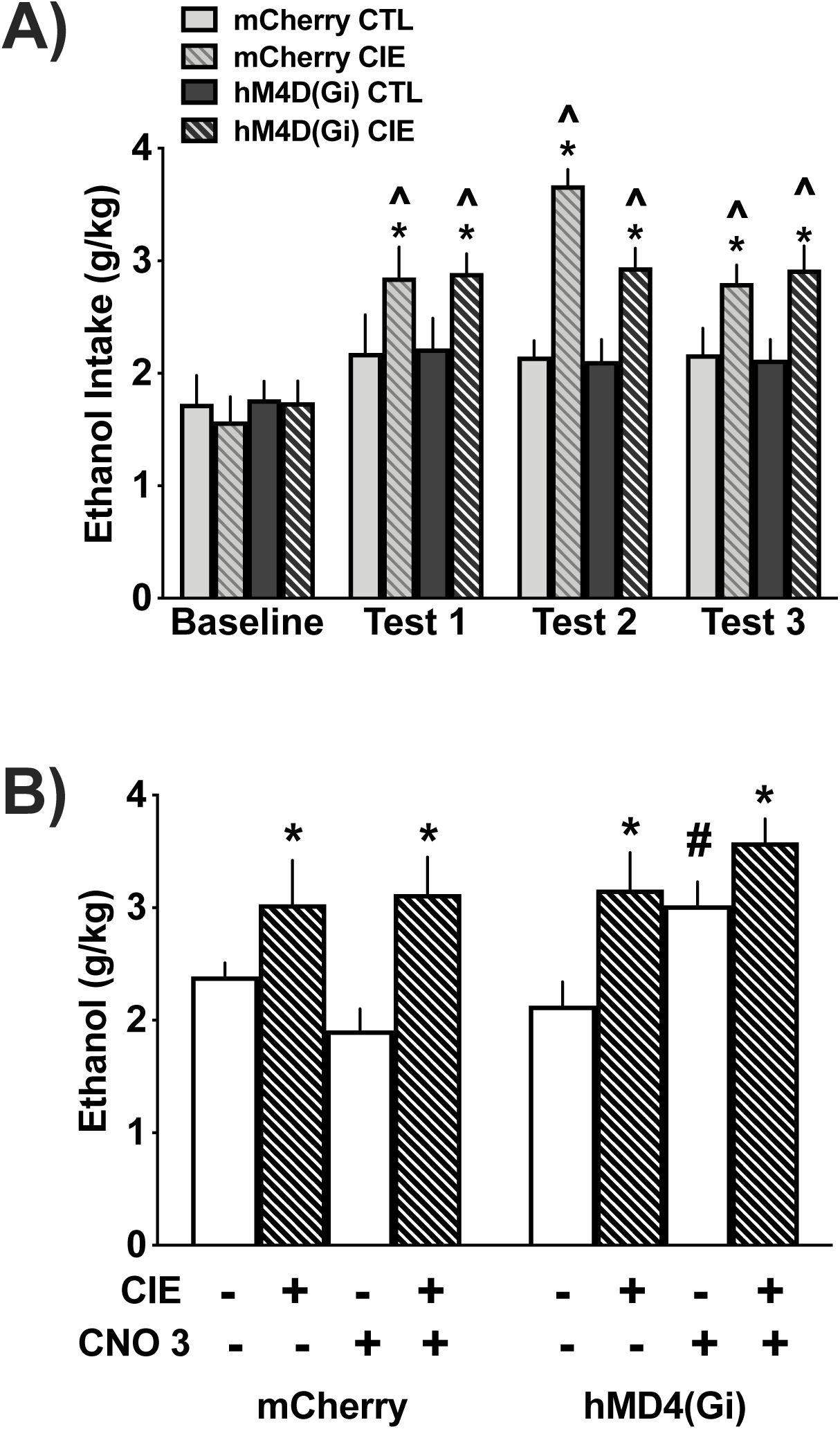
Inhibiting ventral hippocampal function increases drinking in non-dependent (CTL) mice but not in ethanol dependent (CIE) mice. **A)** Ethanol Drinking Prior to CNO Challenge (n=18-20/group). Over time, CIE mice increased ethanol consumption (*p<0.05 within time point, ^p<0.05 versus baseline, regardless of viral construct). **B)** CNO Challenge and Ethanol Drinking (n=8-11/group). CNO had no effects in mice expressing only mCherry (see Results for more details). However, in mice expressing hM4Di in the ventral hippocampus, CNO (3 mg/kg ip.) significantly increased drinking in the CTL mice (#p<0.05 within group for hM4Di virus) without altering the already elevated drinking in the CIE mice. These data suggest that reduced ventral hippocampal function contributes to increased ethanol consumption in ethanol dependence mice. All data are weekly means ± S.E.M.

#### CNO Challenge and Ethanol Drinking

During Test week 4, mice were challenged with CNO in a cross over design on Tuesday and Thursday of that week to activate the inhibitory DREADD, hM4Di, expressed in vHC glutamatergic neurons. These data are summarized in **Figure 1B** and are consistent with the data in Figure 1A showing that CIE mice consume more ethanol than CTL mice. Further, the data show that CNO activation of hM4Di DREADDs in the vHC, i.e., inhibition of vHC projections, only increased ethanol consumption in non-dependent mice. Initially, these data were analyzed using a 3-way ANOVA (2 Groups) X (2 Viruses) X (2 Dose: VEH vs CNO) where Dose was a repeated measure. The 3-way interaction was not significant [F (1,34) =1.7] and neither was the Dose X Group interaction [F (1,34) <0.5]. However, the ANOVA revealed a significant interaction of Virus X Dose [F (1,34) =8.8, p=0.005] indicating that the virus type interacted with CNO challenge.

Additional analyses focused separately on the two different viral groups using a 2-way ANOVA (Group vs Dose), again using Dose as a repeated measure. For the mice expressing only mCherry in the ventral hippocampus, there was no significant interaction [F (1,16) =0.925], nor was there a main effect of Dose [F (1,16) =2.104]. There was, however, a main effect of Group [F (1,16) =11.916, p=0.003], consistent with the increased ethanol consumption by the CIE mice (*p<0.05). Planned comparisons using Sidak’s Post-hoc Test supported the ANOVA indicating significant differences in ethanol consumption between CIE and CTL mice (VEH and CNO conditions both *p<0.05) but did not indicate any other significant differences. The slight decrease in ethanol intake after CNO challenge in the CTL mice was not significant (p=0.09). Thus, as expected, CNO did not alter ethanol consumption in CIE and CTL mice expressing the mCherry control virus.

Similar analyses were conducted on the ethanol consumption data from mice expressing hM4Di in the ventral hippocampus. This analysis did not reveal a significant factor interaction [F (1,18) =0.743] but did indicate significant main effects of Group [F (1,18) =5.996, p=0.025] and Dose [F (1,18) =7.776, p=0.012]. Planned comparisons using Sidak’s Post-hoc Test supported the findings for group differences, but only for the comparison between the CTL and CIE mice after VEH challenge (*p=0.041). No difference was noted between CTL and CIE mice after CNO challenge (p=0.168). While there was a slight increase in ethanol consumption in CIE mice after CNO challenge, this also was not statistically significant (p=0.094). Thus, these data suggest that CNO activation of hM4Di in the vHC increased drinking in CTL mice to amounts observed in CIE mice, but did not significantly affect alcohol intake in the CIE mice.

#### Water Drinking after CNO Challenge

Because water was also available during the ethanol access session, water consumption was also tracked during this study on days when mice were challenged with CNO. Earlier reports from our laboratory using lickometer systems indicate that water consumption is very low in this limited access paradigm (Griffin *et al.* 2009b; Griffin *et al.* 2007). Consistent with these earlier reports, water intake was very low in this experiment. In fact, many values were zero after taking into account the correction for spillage (see methods), though some mice did consume some water. Because water intake was so sparse, no statistical analyses were conducted on these data.

### Experiment 2

#### Ethanol Drinking

All data shown are from ethanol access periods when photometry recording occurred. As expected, the CIE exposure procedures increased ethanol consumption in the CIE group compared to the CTL group (**Figure 2A & 2B**). This was supported by a Group (CIE vs. CTL) X Phase (Baseline vs. Post-CIE) ANOVA with Phase treated as a repeated measure. The factor interaction was significant [F (1,4) =15.076, p =0.018] as was the main effect of Phase [F (1,4) =43.989, p=0.003]. The Group factor was not significant [F (1,4) <2]. Post-hoc analysis indicated that the CIE mice increased drinking during the Post-CIE phase (*p=0.003). Total bottle contacts (per mouse) followed a similar pattern as the ethanol consumption, as would be expected. However, after a similar analysis, there were no significant interaction or main effects found [F (1,4) =1.6]. Lick frequency at the bottle per bout were similar for CTL and CIE mice during both Phases of the study (data not shown).

**Figure 2.**
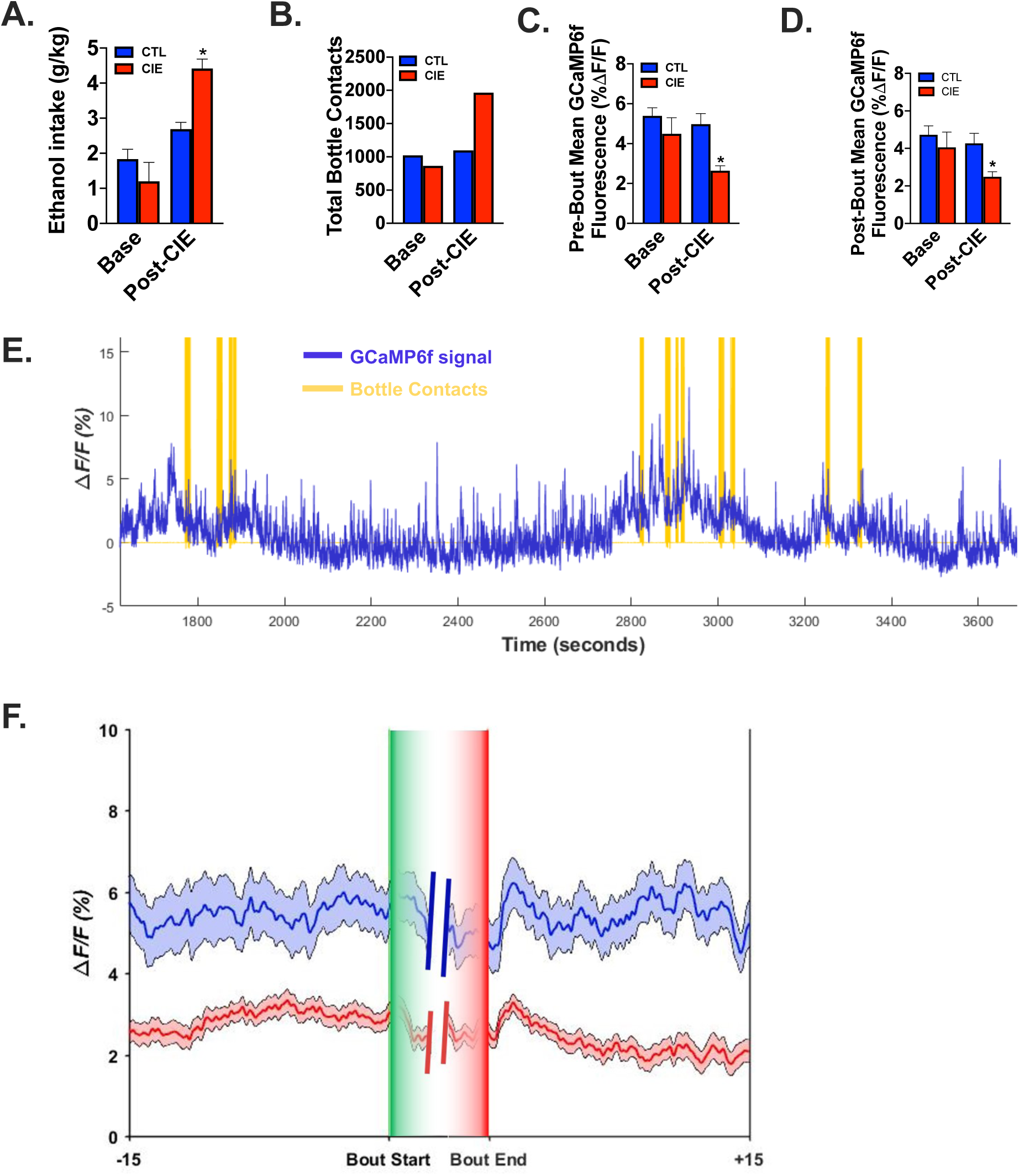
Calcium signaling is attenuated in the ventral hippocampus (vHC) of ethanol dependent (CIE) mice during drinking bouts compared to drinking bouts in non-dependent (CTL) mice (n=3 / group). The data shown in all panels is taken from sessions during which photometry recordings occurred. **A & B)** As expected, CIE exposure increased ethanol consumption (*p<0.05) and total contacts (licks) at the ethanol bottle. **C & D)** The CIE mice showed diminished pre- and post-bout GCaMP6f signals in the vHC compared to the CTL mice (*p<0.05). **E)** An example of calcium signals from a 2-hr recording when mice had access to the bottle containing 15% ethanol. The vertical bars (gold) indicating bottle contacts are actually comprised of many contacts, and in some cases multiple bouts of contacts, ranging from 40-60 contacts per bout. **F)** The normalized calcium signal 15 secs before and after a bout during the Post-CIE phase, the dark line represents the group mean and the shaded area around the line represents standard error of the mean. Data in the bar graphs are means ± S.E.M.

#### Calcium Transients

These analyses focused on bout level events. Here, bouts are defined as contacts occurring over at least a 0.5 sec period at a minimum rate of 6 Hz followed by a 5sec pause before the next bout begins. Along with the consumption data, these data are also summarized in **Figure 2**. In the CTL mice, the normalized pre- and post-bout GCaMP6f signal was elevated prior to and following ethanol drinking bouts, and this signal remained stable over the course of the drinking study. However, after CIE exposure, the CIE mice showed attenuated pre- and post-bout GCaMP6f signals in the ventral hippocampus compared to the CTL mice. The 2-way ANOVA on the pre-bout calcium transients revealed a near-significant factor interaction [F (1,4)=6.456, p=0.064] with a significant main effect of Phase [F (1,4) =6.456, p=0.064]. The group factor was not significant. Post-hoc analysis showed that during the Post-CIE Phase, the CIE mice had reduced pre-bout calcium transients than the CTL mice (**Figure 2C**). Analysis on the post-bout calcium transients revealed similar findings. The factor interaction was significant [F (1,4) =13.353, p=0.022] as was the main effect of Phase [F (1,4) =89.123, p=0.001], while the Group factor was not [F (1,4) =1.362]. Post-hoc analysis showed that during the Post-CIE Phase, the CIE mice had reduced post-bout calcium transients than the CTL mice (**Figure 2D**). Shown in **Figure 2E** is an example recording from a 2-hr drinking period showing the normalized calcium signal (in green) and the bottle contacts in vertical red lines. Note that the vertical red lines obscure the numerous contacts and multiple bouts of ethanol drinking that can occur in relatively short periods of time. Finally, the normalized calcium signal for the 15 secs before and after a bout during the Post-CIE phase are shown in **Figure 2F** with the dark line representing the group mean and the shaded area around the line indicating standard error of the mean. Together, these findings indicate that after mice are rendered ethanol dependent and consume more ethanol, there is less functional activity in the ventral hippocampus surrounding bouts of active ethanol drinking.

## Discussion

The two principle findings of these experiments are that inhibiting the ventral hippocampus increases drinking in non-dependent (CTL) mice, elevating their drinking to levels similar to that of dependent mice, and that ethanol dependent (CIE) mice have attenuated calcium signaling in the ventral hippocampus surrounding drinking bouts. Together, these findings suggest that ethanol dependence reduces activity of the ventral hippocampus and that reduced function of this brain region contributes to the increased drinking phenotype in ethanol dependent mice.

Chemogenetic inhibition of ventral hippocampal activity increased ethanol drinking but, interestingly, this was significant only for the non-dependent mice (CTL group). This suggests that the ventral hippocampus contributes an inhibitory influence on ethanol consumption that is disrupted by ethanol dependence thus facilitating higher ethanol consumption. This is similar to evidence for infralimbic projections to the NAc exerting control over excessive drinking in ethanol dependent rats (Pfarr *et al.* 2015). Prior literature indicates that the hippocampus is part of a behavioral inhibition circuit in the brain that serves to keep certain behaviors in check (McNaughton 2006; Schwarting & Busse 2017). Recent work has explored this concept by modeling approach-avoidance behaviors in rodents. In these studies, data indicate that inhibition of ventral hippocampal circuitry increases approach behaviors to contexts that were previously avoided (Ito & Lee 2016; Schumacher *et al.* 2016). Relevant to the present study are earlier data indicating an inhibitory role of the hippocampus on consummatory behavior (Bannerman *et al.* 2002a). Thus, the findings in this report extend earlier work to include excessive ethanol drinking.

The outcome of the first experiment suggested that reduced function of the ventral hippocampus drives ethanol drinking to higher levels, resembling the phenotype of ethanol dependent mice. To confirm that CIE exposure reduces functional activity in the vHC, we measured calcium transients in neurons with the calcium sensor GCaMP6f expressed the under the control of the CaMKIIa promoter using fiber photometry techniques. Importantly, blunted GCaMP6f signals in the vHC of ethanol dependent mice occurred during a phase of the experiment when they consumed significantly more ethanol than the non-dependent mice. Of course, the blunted GCaMP6f signal in the ventral hippocampus of dependent mice drinking more ethanol is consistent with the chemogenetic experiment showing that inhibition of the vHC increased ethanol drinking in non-dependent mice.

Despite the already reduced activity in the ventral hippocampus of CIE mice shown in the photometry fiber recordings, chemogenetic inhibition of ventral hippocampal activity only produced a small, non-significant additional increase in ethanol drinking in this group. It is possible that the lack of a further increase in drinking by the CIE mice can be attributed to a ceiling effect either caused by the time constraints of the limited access procedure or simply reaching a physiological limit for consumption. However, if the time available for ethanol access is extended, we have found that B6 mice are capable of drinking 6-7 g/kg of 20% ethanol in a 4-hour period (Haun, Lopez, Griffin and Becker, unpublished) indicating that B6 are physiologically capable of drinking more ethanol. In addition, other recent work indicates that B6 mice can consume ∼4.5 g/kg within 2 hours after systemic challenge with the kappa receptor agonist U50488 while drinking 20% ethanol (Haun, Lopez, Griffin and Becker, unpublished). Together, these findings suggest the likely possibility that the 2-hour access period used in these studies may be a constraint on drinking when ethanol dependent mice are drinking 15% ethanol. Alternatively, if providing a longer ethanol access period in combination with chemogenetic inhibition of the vHC in ethanol dependent mice does not further increase consumption, this could suggest a limit on the reduction in vHC activity that will influence drinking. Thus, additional experiments will be required to determine if chemogenetic inhibition of the ventral hippocampus in ethanol dependent mice can increase drinking even further.

In the first experiment, we also measured water intake and found that very little occurs in this limited access drinking procedure, consistent with earlier reports (Griffin *et al.* 2009b; Griffin *et al.* 2007). Despite this limitation, we did not find systematic evidence that chemogenetic inhibition changed consumption of the alternate fluid choice in these mice. Still, it is premature to suggest that the effect of ventral hippocampus inhibition is specific to ethanol. It will be important to investigate the role of the ventral hippocampus in consumption of other reinforcers, especially natural reinforcers like food and sucrose. Some reports indicate the involvement of the ventral hippocampus in models of feeding behavior using high concentrations of sucrose. Data from these experiments show that inhibition of ventral hippocampus function after concluding a meal increases feeding at the next meal (Hannapel *et al.* 2019; Hannapel *et al.* 2017), a finding broadly consistent with the data presented in this report. Thus, more work needs to be done to determine the role of the ventral hippocampus on consummatory behavior.

Human imaging studies indicate that compared to healthy controls there is hippocampal atrophy in patients with alcohol use disorder (Agartz *et al.* 1999; Beresford *et al.* 2006; Nagel *et al.* 2005). While human imaging studies do not differentiate dorsal and ventral areas of the hippocampus, preclinical work in the 1980’s suggests that the ventral portion of the hippocampus may be more susceptible to ethanol-related plasticity, exhibiting larger reductions in spine density (Lescaudron & Verna 1985). Interestingly, more recent work supports the idea that the vHC may be more vulnerable to ethanol exposure with reports demonstrating increased excitability which was associated with reduced expression of key proteins that regulate synaptic transmission in the vHC (Almonte *et al.* 2017; Ewin *et al.* 2019). Although this appears opposite to the data presented here, given the complex interactions between the hippocampal subregions in the ventral hippocampus (CA1, CA2, CA3 and Subiculum), sorting out the multifaceted effects of ethanol on neuronal function in this area and ultimately determining how these alterations influence behavior will be an important undertaking.

In conclusion, reduced function of the ventral hippocampus appears to be permissive in the context of ethanol drinking, leading to increased ethanol consumption. This was measured directly using fiber photometry procedures and also supported by using chemogenetic techniques to inhibit function of this brain region. The work presented here reveals a complexity in hippocampal function with regards to ethanol drinking behavior that requires further study to understand. Much of the work supporting a role for the hippocampus in various behaviors relies on permanent or reversible lesions, however more recent work using newer, more selective technology is emerging that is helping to resolve unanswered questions and, at the same time, reveal the complex function of this heterogenous structure (Barker *et al.* 2019; Britt *et al.* 2012; Ewin *et al.* 2019; Scudder *et al.* 2018; Yu *et al.* 2017). Despite the exciting directions that research in this area is going, many questions regarding ventral hippocampal functional remain unresolved. One important consideration for future efforts is to examine the role of ventral hippocampal connectivity to other limbic areas such as the nucleus accumbens, which is known to be crucial for processing reward-related information. Recent work has identified that this pathway and its engagement of a feed-forward inhibitory circuit within the nucleus accumbens as important to a variety of behaviors (Scudder *et al.* 2018; Yu *et al.* 2017). The role of the vHC to accumbens pathway in ethanol drinking behavior is relatively unexplored but was recently shown to be influenced by ethanol dependence in the same mouse model used in the present studies (Kircher *et al.* 2019). Thus, future work will focus on the ventral hippocampal to accumbens pathway and the role it plays in driving excessive drinking in ethanol dependence.

## Acknowledgements

Technical assistance provided by Sarah K. Brown. Funding provided by R21AA024881 (WCG), P50AA010761 (WCG, HCB, JAR, PJM, JJW), U01AA020930 (PJM), UO1AA014095 (HCB), U24AA020929 (HCB) & VA Medical Research (HCB).

## BIBLIOGRAPHY

Agartz I, Momenan R, Rawlings RR, Kerich MJ, Hommer DW (1999) Hippocampal volume in patients with alcohol dependence. Arch Gen Psychiatry 56: 356–63.

Almonte AG, Ewin SE, Mauterer MI, Morgan JW, Carter ES, Weiner JL (2017) Enhanced ventral hippocampal synaptic transmission and impaired synaptic plasticity in a rodent model of alcohol addiction vulnerability. Sci Rep 7: 12300.

Bannerman DM, Deacon RM, Offen S, Friswell J, Grubb M, Rawlins JN (2002a) Double dissociation of function within the hippocampus: spatial memory and hyponeophagia. Behav Neurosci 116: 884–901.

Bannerman DM, Lemaire M, Yee BK, Iversen SD, Oswald CJ, Good MA, Rawlins JN (2002b) Selective cytotoxic lesions of the retrohippocampal region produce a mild deficit in social recognition memory. Exp Brain Res 142: 395–401.

Bannerman DM, Rawlins JN, McHugh SB, Deacon RM, Yee BK, Bast T, Zhang WN, Pothuizen HH, Feldon J (2004) Regional dissociations within the hippocampus--memory and anxiety. Neurosci Biobehav Rev 28: 273–83.

Barker JM, Bryant KG, Chandler LJ (2019) Inactivation of ventral hippocampus projections promotes sensitivity to changes in contingency. Learn Mem 26: 1–8.

Becker HC (2013) Animal models of excessive alcohol consumption in rodents. Curr Top Behav Neurosci 13: 355–77.

Becker HC, Lopez MF (2004) Increased ethanol drinking after repeated chronic ethanol exposure and withdrawal experience in C57BL/6 mice. Alcohol Clin Exp Res 28: 1829–38.

Beresford TP, Arciniegas DB, Alfers J, Clapp L, Martin B, Beresford HF, D. Y, Liu D, Shen D, Davatzikos C, Laudenslager ML (2006) Hypercortisolism in alcohol dependence and its relation to hippocampal volume loss. J Stud Alcohol 67: 861–7.

Bossert JM, Adhikary S, St Laurent R, Marchant NJ, Wang HL, Morales M, Shaham Y (2016) Role of projections from ventral subiculum to nucleus accumbens shell in context-induced reinstatement of heroin seeking in rats. Psychopharmacology (Berl) 233: 1991–2004.

Braunscheidel KM, Okas MP, Hoffman M, Mulholland PJ, Floresco SB, Woodward JJ (2019) The abused inhalant toluene impairs medial prefrontal cortex activity and risk/reward decision making during a probabilistic discounting task. J Neurosci.

Britt JP, Benaliouad F, McDevitt RA, Stuber GD, Wise RA, Bonci A (2012) Synaptic and behavioral profile of multiple glutamatergic inputs to the nucleus accumbens. Neuron 76: 790–803.

DePoy L, Daut R, Brigman JL, MacPherson K, Crowley N, Gunduz-Cinar O, Pickens CL, Cinar R, Saksida LM, Kunos G, Lovinger DM, Bussey TJ, Camp MC, Holmes A (2013) Chronic alcohol produces neuroadaptations to prime dorsal striatal learning. Proc Natl Acad Sci USA 110: 14783–8.

Dhaher R, Finn D, Snelling C, Hitzemann R (2008) Lesions of the extended amygdala in C57BL/6J mice do not block the intermittent ethanol vapor-induced increase in ethanol consumption. Alcohol Clin Exp Res 32: 197–208.

Ewin SE, Morgan JW, Niere F, McMullen NP, Barth SH, Almonte AG, Raab-Graham KF, Weiner JL (2019) Chronic Intermittent Ethanol Exposure Selectively Increases Synaptic Excitability in the Ventral Domain of the Rat Hippocampus. Neuroscience 398: 144–157.

Fanselow MS, Dong HW (2010) Are the dorsal and ventral hippocampus functionally distinct structures? Neuron 65: 7–19.

Ferbinteanu J, McDonald RJ (2001) Dorsal/ventral hippocampus, fornix, and conditioned place preference. Hippocampus 11: 187–200.

Finn DA, Snelling C, Fretwell AM, Tanchuck MA, Underwood L, Cole M, Crabbe JC, Roberts AJ (2007) Increased Drinking During Withdrawal From Intermittent Ethanol Exposure Is Blocked by the CRF Receptor Antagonist d-Phe-CRF(12-41). Alcohol Clin Exp Res 31: 939–49.

Franklin KBJ, Paxinos G (2008) The Mouse Brain in Stereotaxic Coordinates, Third Edition, 3rd edn. Academic Press, San Diego

Gilpin NW, Richardson HN, Lumeng L, Koob GF (2008) Dependence-induced alcohol drinking by alcohol-preferring (P) rats and outbred Wistar rats. Alcohol Clin Exp Res 32: 1688–96.

Griffin WC, 3rd (2014) Alcohol dependence and free-choice drinking in mice. Alcohol 48: 287–93.

Griffin WC, 3rd, Haun HL, Hazelbaker CL, Ramachandra VS, Becker HC (2014) Increased extracellular glutamate in the nucleus accumbens promotes excessive ethanol drinking in ethanol dependent mice. Neuropsychopharmacology 39: 707–17.

Griffin WC, 3rd, Ramachandra VS, Knackstedt LA, Becker HC (2015) Repeated Cycles of Chronic Intermittent Ethanol Exposure Disrupt Glutamate Homeostasis in Mice. Front Pharmacol 6.

Griffin WC, Lopez MF, Becker HC (2009a) Intensity and duration of chronic ethanol exposure is critical for subsequent escalation of voluntary ethanol drinking in mice. Alcoholism Clin Exp Res 33: 1893–900.

Griffin WC, Lopez MF, Yanke AB, Middaugh LD, Becker HC (2009b) Repeated cycles of chronic intermittent ethanol exposure in mice increases voluntary ethanol drinking and ethanol concentrations in the nucleus accumbens. Psychopharmacology 201: 569–80.

Griffin WC, Middaugh LD, Becker HC (2007) Voluntary ethanol drinking in mice and ethanol concentrations in the nucleus accumbens. Brain Res 1138: 208–13.

Hannapel R, Ramesh J, Ross A, LaLumiere RT, Roseberry AG, Parent MB (2019) Postmeal Optogenetic Inhibition of Dorsal or Ventral Hippocampal Pyramidal Neurons Increases Future Intake. eNeuro 6.

Hannapel RC, Henderson YH, Nalloor R, Vazdarjanova A, Parent MB (2017) Ventral hippocampal neurons inhibit postprandial energy intake. Hippocampus 27: 274–284.

Hansson AC, Rimondini R, Neznanova O, Sommer WH, Heilig M (2008) Neuroplasticity in brain reward circuitry following a history of ethanol dependence. Eur J Neurosci 27: 1912–22.

Ito R, Lee AC (2016) The role of the hippocampus in approach-avoidance conflict decision-making: Evidence from rodent and human studies. Behav Brain Res 313: 345–57.

Jeanes ZM, Buske TR, Morrisett RA (2011) In vivo chronic intermittent ethanol exposure reverses the polarity of synaptic plasticity in the nucleus accumbens shell. J Pharmacol Exp Ther 336: 155–64.

Kanoski SE, Grill HJ (2015) Hippocampus Contributions to Food Intake Control: Mnemonic, Neuroanatomical, and Endocrine Mechanisms. Biol Psychiatry.

Kircher DM, Aziz HC, Mangieri RA, Morrisett RA (2019) Ethanol Experience Enhances Glutamatergic Ventral Hippocampal Inputs to D1 Receptor-Expressing Medium Spiny Neurons in the Nucleus Accumbens Shell. J Neurosci 39: 2459–2469.

Kjelstrup KG, Tuvnes FA, Steffenach HA, Murison R, Moser EI, Moser MB (2002) Reduced fear expression after lesions of the ventral hippocampus. Proc Natl Acad Sci U S A 99: 10825–30.

Lasseter HC, Xie X, Ramirez DR, Fuchs RA (2010) Sub-region specific contribution of the ventral hippocampus to drug context-induced reinstatement of cocaine-seeking behavior in rats. Neuroscience 171: 830–9.

LeGates TA, Kvarta MD, Tooley JR, Francis TC, Lobo MK, Creed MC, Thompson SM (2018) Reward behaviour is regulated by the strength of hippocampus-nucleus accumbens synapses. Nature 564: 258–262.

Lescaudron L, Verna A (1985) Effects of chronic ethanol consumption on pyramidal neurons of the mouse dorsal and ventral hippocampus: a quantitative histological analysis. Exp Brain Res 58: 362–7.

Lopez MF, Becker HC (2005) Effect of pattern and number of chronic ethanol exposures on subsequent voluntary ethanol intake in C57BL/6J mice. Psychopharmacology 181: 688–96.

Marchant NJ, Campbell EJ, Whitaker LR, Harvey BK, Kaganovsky K, Adhikary S, Hope BT, Heins RC, Prisinzano TE, Vardy E, Bonci A, Bossert JM, Shaham Y (2016) Role of Ventral Subiculum in Context-Induced Relapse to Alcohol Seeking after Punishment-Imposed Abstinence. J Neurosci 36: 3281–94.

Maren S, Holt WG (2004) Hippocampus and Pavlovian fear conditioning in rats: muscimol infusions into the ventral, but not dorsal, hippocampus impair the acquisition of conditional freezing to an auditory conditional stimulus. Behav Neurosci 118: 97–110.

McNaughton N (2006) The role of the subiculum within the behavioural inhibition system. Behav Brain Res 174: 232–50.

Nagel BJ, Schweinsburg AD, Phan V, Tapert SF (2005) Reduced hippocampal volume among adolescents with alcohol use disorders without psychiatric comorbidity. Psychiatry Res 139: 181–90.

O’Dell LE, Roberts AJ, Smith RT, Koob GF (2004) Enhanced alcohol self-administration after intermittent versus continuous alcohol vapor exposure. Alcohol Clin Exp Res 28: 1676–82.

Pennartz CM, Ito R, Verschure PF, Battaglia FP, Robbins TW (2011) The hippocampal-striatal axis in learning, prediction and goal-directed behavior. Trends Neurosci 34: 548–59.

Pfarr S, Meinhardt MW, Klee ML, Hansson AC, Vengeliene V, Schonig K, Bartsch D, Hope BT, Spanagel R, Sommer WH (2015) Losing Control: Excessive Alcohol Seeking after Selective Inactivation of Cue-Responsive Neurons in the Infralimbic Cortex. J Neurosci 35: 10750–61.

Rehm J, Dawson D, Frick U, Gmel G, Roerecke M, Shield KD, Grant B (2014) Burden of disease associated with alcohol use disorders in the United States. Alcohol Clin Exp Res 38: 1068–77.

Rehm J, Gmel GE, Sr., Gmel G, Hasan OSM, Imtiaz S, Popova S, Probst C, Roerecke M, Room R, Samokhvalov AV, Shield KD, Shuper PA (2017) The relationship between different dimensions of alcohol use and the burden of disease-an update. Addiction 112: 968–1001.

Rinker JA, Gioia D, Braunscheidel KM, Wayman WN, Hoffman M, Passarella L, Calipari E, Mulholland PJ, Woodward JJ (2018) Monitoring Neural Activity During Exposure to Drugs of Abuse With In Vivo Fiber Photometry. BioRxiv https://doi.org/10.1101/487546.

Roberts AJ, Cole M, Koob GF (1996) Intra-amygdala muscimol decreases operant ethanol self-administration in dependent rats. Alcohol Clin Exp Res 20: 1289–98.

Rogers JL, See RE (2007) Selective inactivation of the ventral hippocampus attenuates cue-induced and cocaine-primed reinstatement of drug-seeking in rats. Neurobiol Learn Mem 87: 688–92.

Schumacher A, Vlassov E, Ito R (2016) The ventral hippocampus, but not the dorsal hippocampus is critical for learned approach-avoidance decision making. Hippocampus 26: 530–42.

Schwarting RK, Busse S (2017) Behavioral facilitation after hippocampal lesion: A review. Behav Brain Res 317: 401–414.

Scudder SL, Baimel C, Macdonald EE, Carter AG (2018) Hippocampal-Evoked Feedforward Inhibition in the Nucleus Accumbens. J Neurosci 38: 9091–9104.

Strange BA, Witter MP, Lein ES, Moser EI (2014) Functional organization of the hippocampal longitudinal axis. Nat Rev Neurosci 15: 655–69.

Sun W, Rebec GV (2003) Lidocaine inactivation of ventral subiculum attenuates cocaine-seeking behavior in rats. J Neurosci 23: 10258–64.

Yu J, Yan Y, Li KL, Wang Y, Huang YH, Urban NN, Nestler EJ, Schluter OM, Dong Y (2017) Nucleus accumbens feedforward inhibition circuit promotes cocaine self-administration. Proc Natl Acad Sci U S A 114: E8750–E8759.

